# H-NS is a conserved repressor of the type VI secretion system in *Vibrio fischeri*

**DOI:** 10.1101/2025.01.30.634523

**Authors:** Lauren Speare, Morgan Pavelsky, Aundre Jackson, Alecia N. Septer

## Abstract

The type VI secretion system (T6SS) is a broadly distributed interbacterial weapon found in both beneficial and pathogenic bacteria and can enhance a microbe’s ability to colonize a host. *Vibrio fischeri* is a beneficial symbiont of fish and squid and a model organism for T6SS function, which is activated in high-viscosity conditions. Previously, we isolated an *hns* mutant in a transposon screen to identify regulators of the T6SS in the fish symbiont *V. fischeri* MJ11. The *hns* gene encodes the DNA-binding protein, H-NS, a conserved global regulator of gene expression that aids in adaptation to changing environments. Quantitative transcriptomes of the *hns* mutant and parent strains grown in liquid or hydrogel media revealed *hns* is required for the global transcriptional changes that occur during transition from lower to higher viscosity conditions. Furthermore, T6SS gene transcripts are more abundant in the *hns* mutant in both conditions, suggesting H-NS represses T6SS in the parent. Single-cell fluorescence microscopy confirmed *hns* mutant cells make more T6SS weapons in both liquid and hydrogel medium, where the *hns* mutant is more proficient at killing a competitor strain, compared to the wild-type parent. Finally, disrupting the *hns* gene in additional light organ isolates resulted in a similar derepression of T6SS, indicating H-NS is a conserved repressor of this interbacterial weapon. This work furthers our understanding of the role of H-NS as a global regulator during environmental shifts in a host-associated bacterial symbiont and expands the list of species where H-NS represses T6SS to include *V. fischeri*.

## Importance

The type VI secretion system (T6SS) is a contact-dependent interbacterial weapon used to eliminate competitors of a niche. Each armed cell contains from one to over six nanoweapons, depending on environmental conditions. Because each weapon is composed of thousands of protein subunits, bacteria must strike a balance between expending metabolic energy on building and deploying weapons and other essential cellular functions, including growth. Here, we show that a single global regulator of gene expression, H-NS, represses synthesis of this weapon under conditions when contact with competitors is low, and prevents cells from becoming too heavily armed under conditions favoring bacterial battles. Although reducing the number of weapons per cell may appear to be a counter-intuitive strategy for defeating a competitor, a similar tactic has been proposed for human conflicts, where a larger number of moderately armed individuals is expected to outcompete a smaller group of more heavily armed individuals.

## Results

The type VI secretion system (T6SS) is an interbacterial weapon found in 25% of Gram-negative bacterial genomes (1), and can be used by the beneficial symbiont, *Vibrio fischeri* for interstrain competition during colonization of its squid host (2, 3). Previously, we performed a transposon mutant screen in the fish symbiont, *V. fischeri* MJ11, to identify genes required for T6SS-mediated killing in hydrogel, a medium that mimics the physical environment within host mucus (4). During this screen, we isolated a mutant (LAS35E11) with a transposon insertion in VFMJ11_1751 (5), which encodes for the broadly-conserved H-NS protein. H-NS is a DNA-binding protein that has global regulatory effects (6), including regulation of symbiosis factors in *V. fischeri* (5, 7, 8), and T6SS activity in other species (9-13). Although this mutant was still able to kill target cells and was therefore a false positive in the previous screen, we were interested in exploring the role of H-NS in regulating gene expression changes, including T6SS, in *V. fischeri*. Therefore, we used a combination of transcriptomics, single-cell microscopy, and coculture assays, to investigate the role of H-NS in regulating global gene expression in *V. fischeri*, and its impact on T6SS.

### H-NS is required for global transcriptional changes during simulated habitat transition

Given that we have previously observed a large transcriptional upshift when wild-type *V. fischeri* cells are grown in higher viscosity medium (4), we wanted to uncover the role H-NS plays in mediating this environmental adaptation by using quantitative transcriptomics, as described previously (4).

We first assessed the transcriptional differences for each of four treatments: wild-type or *hns* mutant grown in liquid or hydrogel (high viscosity) media. We performed a principal coordinate analysis (PCA) using the quantitative transcriptome data and found that the wild-type cultures grown in hydrogel formed a distinct cluster apart from the other three treatments (Fig 1A). To identify the transcriptional changes driving this separation, we created a heatmap that assigns a color for relative transcript abundance for each gene across treatments. This analysis clearly showed the transcriptional upshift we previously reported for WT in hydrogel (4), and that the *hns* mutant was unable to mediate this upshift (Fig 1B). Taken together, these findings suggest that H-NS is required for *V. fischeri* to properly modulate gene expression upon transition from low to high viscosity conditions.

**Figure 1.**
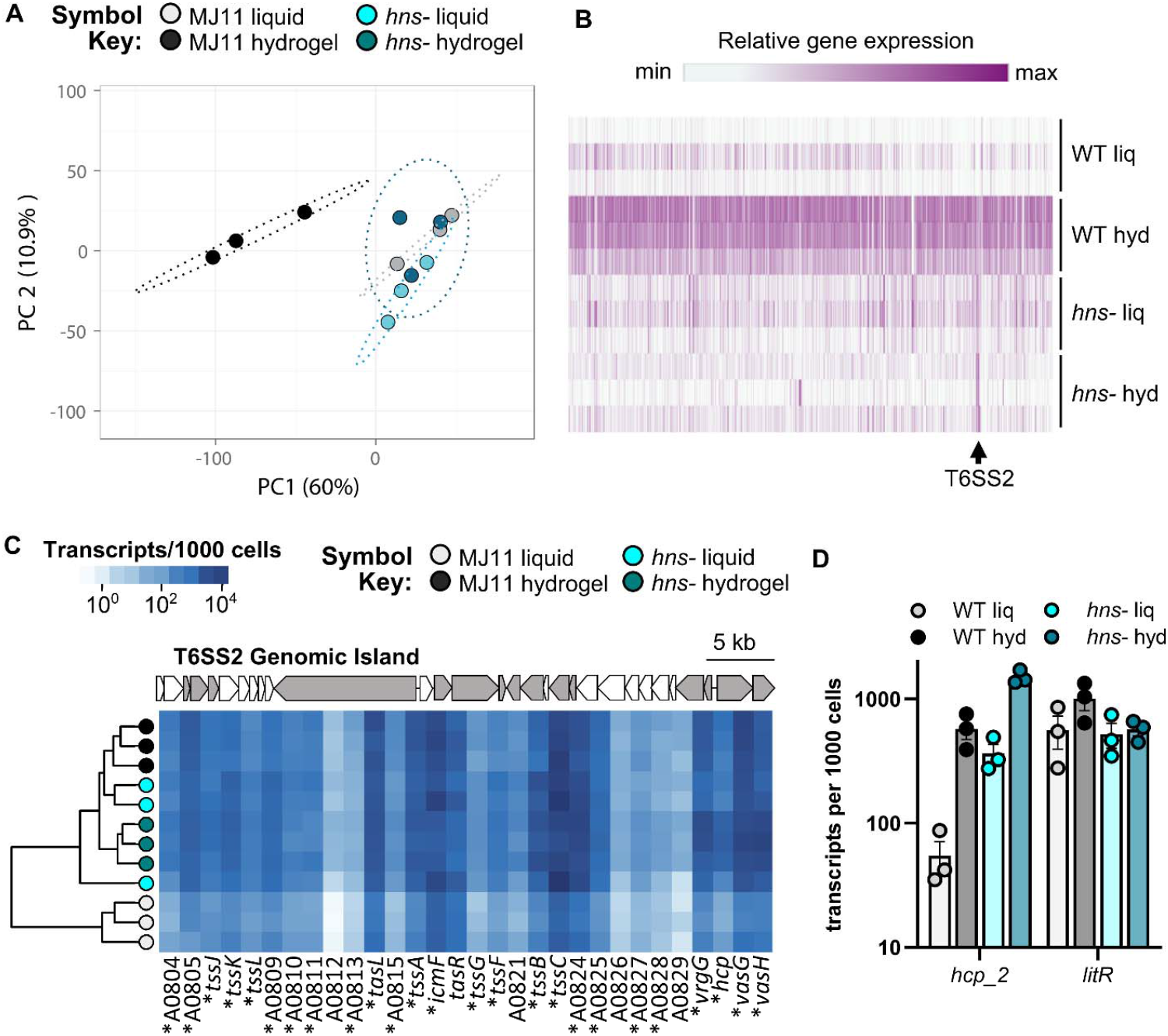
H-NS is necessary for gene expression changes during habitat transition. (A) PCA plot using ClustVis of entire transcriptomes. Unit variance scaling is applied to rows; SVD with imputation is used to calculate principal components. X and Y axis show principal component 1 and principal component 2 that explain 60% and 10.9% of the total variance, respectively. Ellipses indicate 95% confidence. (B) Relative abundance heat map for all genes across four, triplicate treatments. Shading indicates relative change in expression for each gene across treatments with white being the minimum expression level and dark magenta the max. Made with Morpheus from The Broad. (C) Heatmap of hierarchical clustering results for the T6SS2 gene cluster (VFMJ11_A0804-A0833) indicating transcripts per 1000 cell for MJ11 wild-type (WT) grown in liquid (gray) or hydrogel (black) and *hns* mutant grown in liquid (cyan) in liquid or hydrogel (dark cyan). Each row represents a sample and each column represents a gene; gene ID is shown at the bottom of the lower heatmap in each panel. Square color in the heatmap indicates the absolute abundance of each transcript per cell. Asterisks indicate statistically significant differences comparing WT and *hns-* in liquid (t-test, p<0.05). D) Transcripts per 1000 cells for *hcp_2* and *litR* genes for all four treatments. Error bars indicate SEM.

### The T6SS2 gene cluster is highly expressed in *hns* mutant liquid cultures

When exploring our transcriptomes, we noticed the T6SS gene cluster on chromosome II (T6SS2) appeared to be highly expressed in the *hns* mutant, independent of growth condition (Fig 1B). To further explore this observation, we performed a hierarchical clustering analysis using the absolute abundance transcript values for each gene in the T6SS2 gene cluster (Fig 1C). This analysis revealed several important findings. First, the transcript abundance for each T6SS2 gene showed wide variation, spanning four orders of magnitude, depending on the encoded subunit. This observation is consistent with previous work that estimates each T6SS sheath may be comprised of thousands of sheath (TssBC) and inner tube (Hcp) components (14). Second, the T6SS2 transcriptional profiles for both *hns* mutant cultures clustered with the wildtype hydrogel profile, indicating they are most similar and distinct from the wildtype in liquid. Finally, transcript abundance for *litR*, a recently identified negative regulator of T6SS2 in strain FQ-A001 (15), remained high and unchanged across treatments and did not negatively correlate with *hcp* transcript abundance (Fig 1D), suggesting that H-NS mediated repression of T6SS2 is independent of *litR* transcript levels in strain MJ11. These data led us to hypothesize that T6SS2 may be derepressed in liquid-grown *hns* mutant cells, conditions where few sheaths per cell are normally observed in wildtype (16).

### The *hns* mutant makes more T6SS sheaths

To quantify T6SS2 sheath production in the *hns* mutant we moved a TssB/VipA-GFP expression vector (2) into the *hns*::tn5 mutant and grew wild-type or *hns* mutant cultures in liquid or hydrogel media supplemented with IPTG to induce expression of the GFP-tagged sheath protein. Cultures were subsampled at an OD of 1.0, vortexed to disrupt aggregates, and spotted onto glass slides to obtain images of sheaths. For each field of view the number of sheaths per cell was quantified and presented as a percentage of the cells with 0, 1, 2, or 3 or more sheaths per cell. Representative images of wildtype and *hns* mutant cultures grown in liquid and hydrogel are shown in Fig 2A and 2B, respectively. Consistent with our transcriptome data, the proportion of cells with sheaths was higher for the *hns* mutant, compared to the wildtype parent, for both liquid and hydrogel conditions (Fig 2C). The *hns* mutant cells also contained more sheaths per cell, compared to the wildtype (Fig 2D). In combination with our transcriptome data, these findings indicate that H-NS represses T6SS2 gene expression in liquid and suppresses sheath synthesis in both liquid and hydrogel conditions, underscoring the importance of directly visualizing sheaths to fully understand the physiological impact of transcriptional changes.

**Figure 2.**
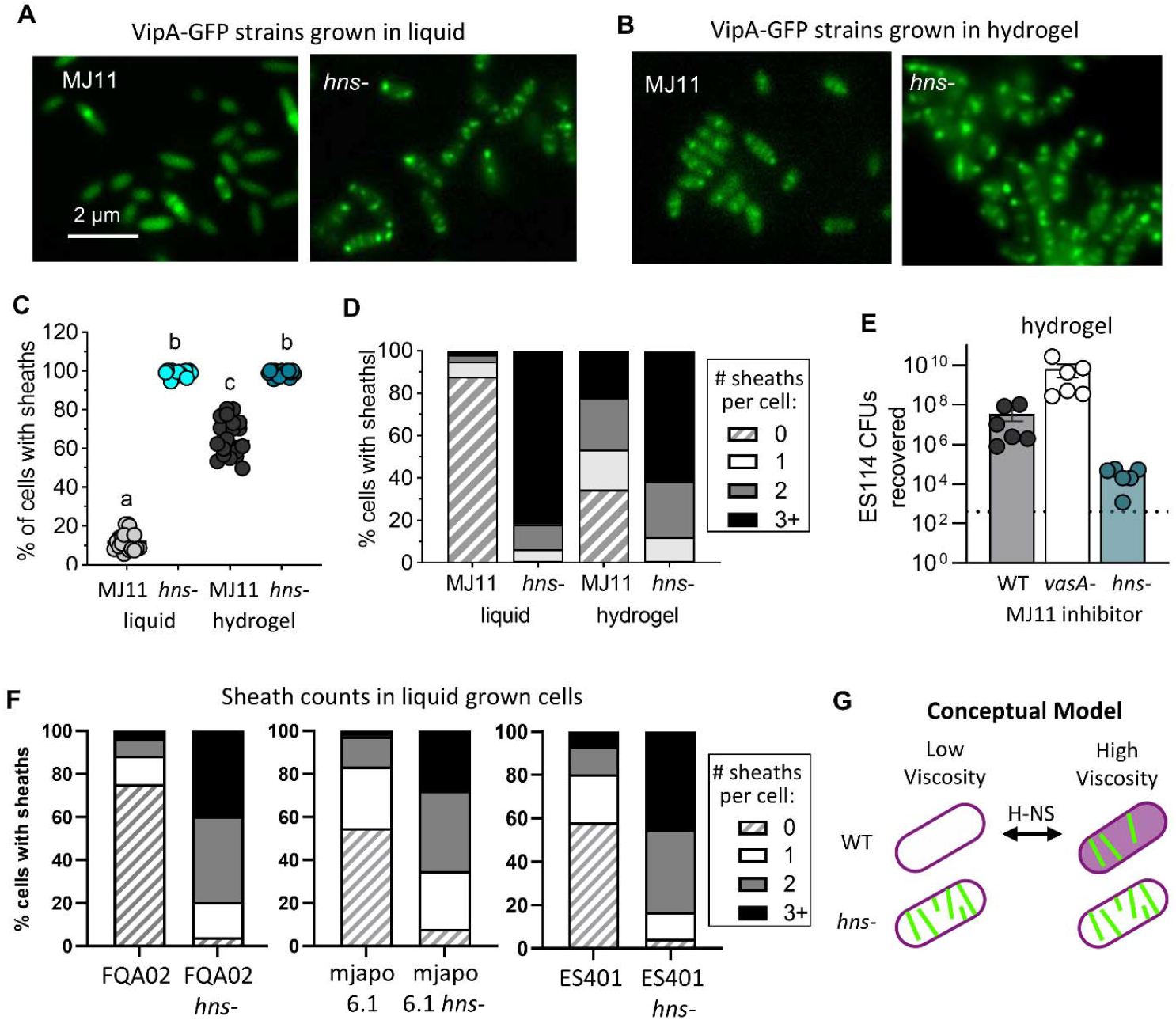
H-NS-mediated repression of T6SS sheaths in liquid is conserved and enhances killing in hydrogel. (A-B) Representative fluorescence microscopy images of wild-type (WT) and *hns-* strains harboring an IPTG-inducible VipA_2-GFP expression vector incubated in either liquid (A) or hydrogel (B) media supplemented with 1.0 mM IPTG for two hours; scale bar = 2 µm. (C) Percentage of cells that contained at least one sheath after two hours in either liquid or hydrogel medium supplemented with 0.5 mM IPTG for two hours. Letters indicate a significantly different percentage of cells with sheaths between treatments. (D) Stacked bar graph showing the percentage of cells within a sample with O (hashed), 1 (white), 2 (light gray), or 3 or more (dark gray) sheaths per cell for MJ11 wild-type and *hns-* cells incubated in liquid or hydrogel medium supplemented with 0.5 mM IPTG for two hours. Each experiment was performed twice with two biological replicates and five fields of view per replicate {n=20). (E) Recovery of ES114 pVSV102 target CFUs after 24 h coincubations with indicated MJ11 strain in 2 ml hydrogel medium in 12 well plate at room temperature. Data shown are combined from two independent experiments, each with three biological replicates. (F) Sheath data for ES401, FQA002, and mjapo6.1 and their hns::tn5 mutants grown in LBS with 0.5 mM IPTG to an OD of −1.0. Sheath percentages based on >400 cells across multiple fields of view. (G) Conceptual model for HNS regulation in *V. fischeri* during environmental change. Cell shading indicates degree of global transcriptional change (white, low; magenta, high), green lines represent T6SS2 sheaths.

Given that *hns* mutants are more heavily armed than their wildtype parents, we asked how this might impact competitor elimination. We performed coincubation assays with wildtype or *hns* mutant inhibitor strains with the ES114 target strain in hydrogel and found that fewer ES114 target CFUs were recovered when coincubated with the *hns* mutant, compared to the wildtype parent (Fig 2E), suggesting heavily armed cells can more effectively reduce competitor numbers in coculture.

### H-NS is a conserved repressor of T6SS2 in *V. fischeri*

We next asked whether H-NS repression of T6SS2 is conserved across diverse *V. fischeri* isolates. To answer this question, we used natural transformation to move the *hns*::tn5 mutation from LAS35E11 into strains ES401, FQ-A002, and mjapo6.1. The TssB/VipA-GFP expression vector was moved into each parent and *hns* mutant strain, and sheath images were quantified for cultures grown in liquid LBS. We observed derepression of T6SS sheath synthesis for all three additional *hns* mutant strains tested (Fig 2F), indicating H-NS represses T6SS broadly in *V. fischeri*.

In summary, the *V. fischeri hns* mutant was impaired in modulating gene expression changes that occur in wildtype upon transition from low to high viscosity conditions (Fig 2G). Furthermore, *hns* mutants are derepressed for T6SS2 gene expression and sheath formation, resulting in more armed cells within a population and more sheaths per cell. These heavily armed populations more efficiently eliminate unarmed target cells in competition. What remains unknown is the mechanism of H-NS repression in wild-type cells and the benefit of limiting the degree to which a population is armed. One might argue that having more heavily armed cells is a better competitive strategy. However, given the predicted metabolic cost to building and using the T6SS2 in *V. fischeri* (17), H-NS-mediated suppression of T6SS2 may be an evolutionary strategy to strike a balance between using energy for cell growth vs arming cells. Indeed, Lanchester’s square law argues that for certain human conflicts, having a higher number of less armed individuals can be more beneficial than fewer, heavily armed individuals (18). Perhaps *V. fischeri*, and other bacteria with H-NS repression of T6SSs, have evolved a similar strategy to increase the chances of a favorable outcome in battle.

## Methods

See supplementary methods document for details on strains and plasmids, quantitative transcriptomes, sheath imaging, and coincubation assays.

## Data Availability

Transcriptome data are available at GenBank under BioProject ID PRJNA1013100.

## Acknowledgements

Work in the Septer lab was supported by NIH NIGMS grant R35 GM137886 and Gordon and Betty Moore Foundation grant GBMF9328. LS was supported by a UNC dissertation completion fellowship.

