## Supplemental methods for "H-NS is a conserved repressor of the type VI secretion system in *Vibrio fischeri*"

**Supplemental methods for Speare et al.:**

**Bacterial strains, plasmids, and media.** *V. fischeri* strains were cultivated at 24-5°C in Luria-Bertani salt (LBS) medium (per liter: 10 g Tryptone, 5 grams Yeast Extract, 20 g NaCl, 50 mL 1 M Tris buffer [pH 7.5]). When indicated, LBS was supplemented with 5% w/vol polyvinylpyrrolidone (PVP360, Sigma-Aldrich) to make a hydrogel medium. *E. coli* strains used for cloning and were grown at 37°C in Luria-Bertani (LB) medium (per liter: 10 g Tryptone, 5 g Yeast Extract, 10 g NaCl) or Brain Heart Infusion (BHI). Media was supplemented with 15 g per liter agar for plates. Antibiotics were added at the following final concentrations in the media: kanamycin, 100 μg/mL for *V. fischeri* and 50 μg/mL for *E. coli*; erythromycin, 5 μg/mL for *V. fischeri* and 150 μg/mL for *E. coli* grown in BHI medium. Bacterial strains and plasmids are listed in the table below.

| **Strain** | **Relevant characteristics** | **Reference** |
| --- | --- | --- |
| ES114 | *V. fischeri;* isolated from *Euprymna scolopes* light organ | Boettcher and Ruby 1990 |
| ES401 | *V. fischeri;* isolated from *Euprymna scolopes* light organ | Fidiopastis *et al*., 2002 |
| MJ11 | *V. fischeri;* isolated from *Moncentris japonica* light organ | Ruby and Nealson 1976 |
| LAS35E11 | MJ11 *hns::tn5* mutant (Erm^R^) | Zarate *et al*., 2024 |
| MP110 | ES401 *hns::tn5* made from LAS35E11 gDNA (Erm^R^) | Zarate *et al*., 2024 |
| FQA002 | *V. fischeri;* isolated from *Euprymna scolopes* light organ | Speare *et al*., 2018 |
| MP108 | FQA002 *hns::tn5* made from LAS35E11 gDNA (Erm^R^) | This study |
| mjapo6.1 | *V. fischeri;* isolated from *Moncentris japonica* light organ | Mandel *et al*., 2009 |
| MP107 | mjapo6.1 *hns::tn5* made from LAS35E11 gDNA (Erm^R^) | This study |
| *E. coli* CC118λpir | *E. coli; Δ(ara-leu) araD Δlac74 galE galK phoA20 thi-1 rpsE rpsB argE*(Am) *recA λpir* | Herrero *et al.,* 1990 |
| *E. coli* DH5⍺λpir | *λpir derivative of E. coli; F’/endA1 hsdR17 glnV44 thi-1 recA1 gyrA relA1* Δ*(lacIZYAargF)*  *U169deoR(f80dlacI* Δ(*lacZ)M15)* | Dunn *et al.,* 2005 |
| **Plasmids** | **Relevant characteristics** | **Reference** |
| pEVS104 | conjugative helper, *oriV_R6k_*_γ_, *oriT, Kn^R^* | Stabb and Ruby 2002 |
| pLostfox-Kn | *tfoX* expression vector, *oriT*, *f1 ori*, *Kn*^R^ | Brooks *et al.,* 2014 |
| pVSV102 | GFP expression vector*, oriV_R6k_*_γ_, *oriV_pES213_, oriT, Kn^R^* | Dunn *et al*., 2006 |
| pSNS119 | IPTG-indicuble vipA-GFP expression vector*, oriV_R6k_*_γ_, *oriV_pES213_, oriT, Kn^R^* | Speare *et al*., 2018 |

**Transcriptomes.** RNA was collected and transcriptomes sequenced for MJ11 WT and the LAS35E11 *hns*::tn5 mutant grown in liquid LBS (liq) or hydrogel, LBS with 5% w/v polyvinylpyrrolidone (PVP) as described in Speare et al., 2024. Transcripts per 1000 cells for all genes are provided in Table S1 of Zarate et al., 2024, and raw reads can be found in GenBank under BioProject ID PRJNA1013100.

**Natural transformation.** To move the *hns*::tn5 mutation from MJ11 strain LAS35E11 into other *V. fischeri* isolates, genomic DNA was isolated from LAS35E11 using a Zymo Quick-DNA Fungal/Bacterial Miniprep Kit. Recipient strains FQA002 and mjapo6.1 were transformed with pLostfox-Kn and cells were prepared for natural transformation as described in (Brooks et al., 2014). Transformed cells were plated onto LBS Erm to select for the mutation of interest (*hns*::tn5), which encode erythromycin resistance. Individual colonies were picked, restreaked and verified to have acquired the mutation (Erm-resistant) and lost the plasmid (Kan-sensitive).

**Coincubation assays.** Killing ability of MJ11-derived strains were assayed by mixing OD 1.0 cell suspensions (grown on overnight LBS plates at 24°C) of MJ11 strain with GFP-tagged ES114 target at a 9:1 ratio and diluting the mixture 100-fold into 2 ml LBS hydrogel (LBS + 5% w/v polyvinylpyrrolidone (PVP)) in 12 well plates that were incubated at room temperature for 24 hours. ES114 pVSV102 target CFUs were recovered by plating serial dilutions of coincubations onto LBS Kan, to select for ES114 pVSV102.

**Single cell fluorescence microscopy and sheath quantification**. To image sheaths in indicated strains, triparental mating were used with the *V. fischeri* recipient strain, pEVS104 conjugative helper, and *E. coli* donor strain carrying the IPTG-inducible VipA/TssB-GFP expression vector. To image and quantify sheaths, pSNS119-containing strains were grown in indicated media (liquid LBS or LBS +PVP) supplemented with 0.5 mM IPTG for 2 hr (Fig 2A-D) or to an OD of 1.0 (Fig 2F) and spotted onto a glass slide to capture green fluorescence images using an Olympus BX51 microscope outfitted with a Hammatsu C8484-03G01 camera (Fig 2A-D) or a Nikon Ti2 inverted fluorescence microscope equipped with a Hamamatsu Orca Fusion camera and NIS Elements software (Fig 2F). Each treatment had at least two biological replicates and images captured across at least five fields of view to capture a minimum of 400 cells. Sheath counts were done manually.
